# Modeling airway dysfunction in asthma using synthetic mucus biomaterials

**DOI:** 10.1101/2020.10.16.342766

**Authors:** Daniel Song, Ethan Iverson, Logan Kaler, Shahed Bader, Margaret A. Scull, Gregg A. Duncan

**Affiliations:** Fischell Department of Bioengineering, University of Maryland, College Park, MD 20742, USA; Department of Cell Biology & Molecular Genetics, and University of Maryland, College Park, MD 20742, USA; Biophysics Program, University of Maryland, College Park, MD 20742, USA

**Keywords:** respiratory disease, mucus, biomaterials, asthma, influenza

## Abstract

As asthma worsens, occlusion of airways with mucus significantly contributes to airflow obstruction and reduced lung function. Recent evidence from clinical studies has shown mucus obtained from adults and children with asthma possesses altered mucin composition. However, how these changes alter the functional properties of the mucus gel is not yet fully understood. To study this, we have engineered a synthetic mucus biomaterial to closely mimic the properties of native mucus in health and disease. We demonstrate this model possesses comparable biophysical and transport properties to native mucus *ex vivo* collected from human subjects and *in vitro* isolated from human airway epithelial (HAE) tissue cultures. We found by systematically varying mucin composition that mucus gel viscoelasticity is enhanced when predominantly composed of mucin 5AC (MUC5AC), as is observed in asthma. As a result, asthma-like synthetic mucus gels are more slowly transported on the surface of HAE tissue cultures and at a similar rate to native mucus produced by HAE cultures stimulated with the type 2 cytokine IL-13, known to contribute to airway inflammation and MUC5AC hypersecretion in asthma. We also discovered the barrier function of asthma-like synthetic mucus towards influenza A virus was impaired as evidenced by the increased frequency of infection in MUC5AC-rich hydrogel coated HAE cultures. Together, this work establishes a biomaterial-based approach to understand airway dysfunction in asthma and related muco-obstructive lung diseases.

**Graphical Abstract:** 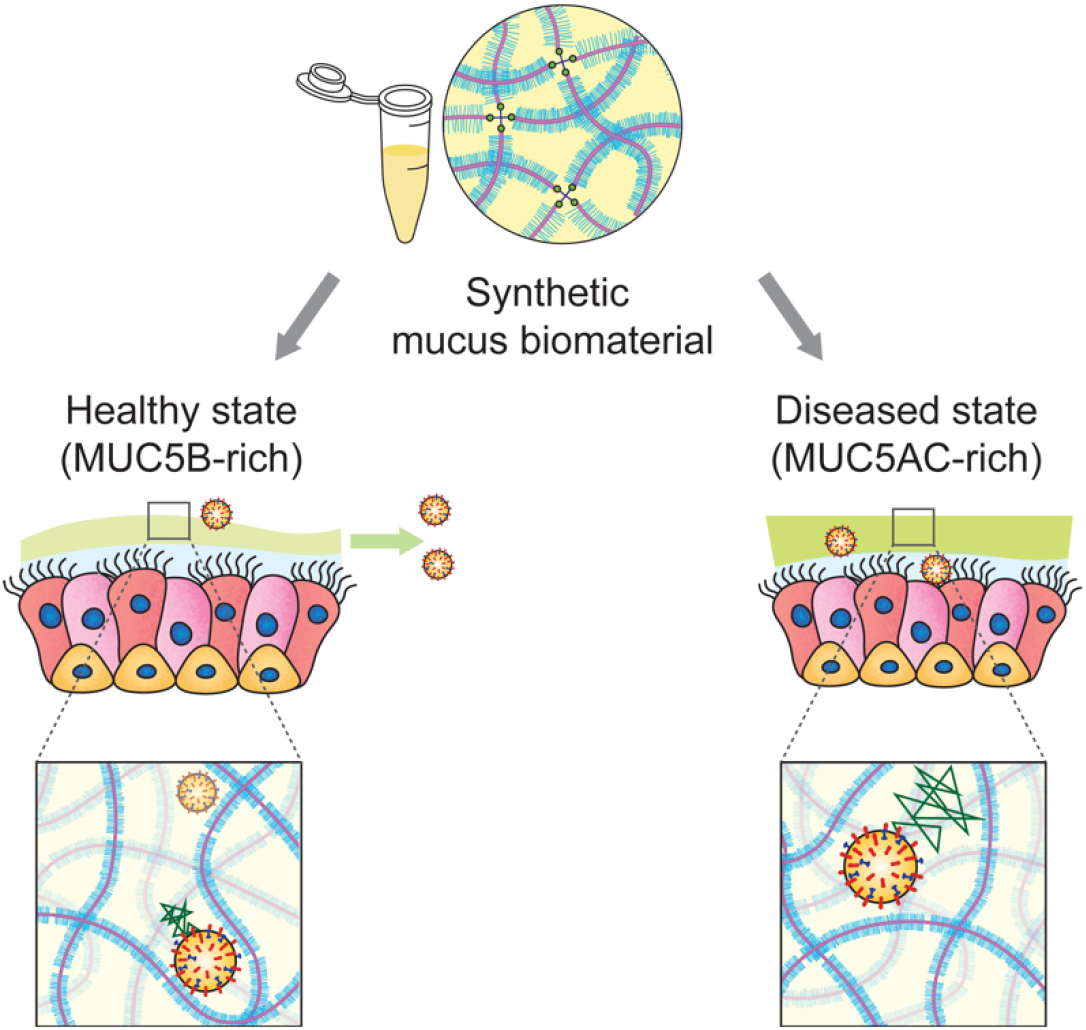

## 1. Introduction

Asthma affects ~300 million people worldwide of all racial and ethnic backgrounds with cases documented from childhood to adulthood [1]. As lung disease worsens in asthma, chronic airway inflammation commonly leads to excess production of mucus [2,3]. Mucus is continuously cleared from the lung through high frequency, coordinated beating of ciliated cells on the surface of the airway epithelium [4,5]. This process, known as mucociliary clearance (MCC), provides the lung with an important defense mechanism against infection and injury with the continuous removal of inhaled environmental particles captured within the mucus layer [6]. However, in all stages of asthma, MCC mechanisms become impaired, resulting in the accumulation of mucus in the airway [7]. Persistent mucus accumulation not only leads to inflammation and infection by providing an environment for microbial growth, but also results in obstruction of conducting airways potentially causing asphyxiation in severe cases of asthma [8,9]. Further, many individuals with asthma suffer from recurrent pulmonary exacerbations characterized as an acute worsening of symptoms requiring emergency department visits and hospitalizations [10,11]. Exacerbations in asthma are often associated with infections by respiratory viruses such as rhinovirus, respiratory syncytial virus (RSV), and influenza [12–14] Accordingly, it is of importance to understand how the changes to mucus produced in asthma leads to dysfunctional MCC and further examine its potential role in susceptibility to infection.

The primary building blocks of respiratory mucus are the gel-forming mucins, mucin 5B (MUC5B) and mucin 5AC (MUC5AC) [15]. These large (~MDa) and heavily glycosylated proteins form the mucus gel network primarily through reversible associations (e.g. entanglements, hydrogen bonds) and disulfide crosslinks [15]. In individuals with asthma, biochemical analysis of mucus produced by cough has revealed mucin composition is altered as a function of disease severity, with a shift from MUC5B to MUC5AC as the predominant secreted mucin [16]. Further, computerized tomography (CT) imaging has revealed an increased likelihood and number of mucus plugs within the airway in asthma patients with MUC5AC-enriched airway mucus [17] These data suggest overproduction of MUC5AC may predispose the airways to mucus accumulation in asthma. MCC defects in asthma have been attributed to oxidation-induced crosslinking of mucins in the asthmatic lung and physical tethering of MUC5AC to the airway surface [18–20], both as a result of exaggerated inflammatory responses in asthma. However, it is not yet fully understood how an imbalance in the ratio of MUC5B to MUC5AC contributes to the biological function of mucus in asthma.

Studying airway mucus in health and disease has relied heavily on acquiring *ex vivo* human mucus samples through sputum induction or endotracheal tubes from patients who have undergone intubation [21,22]. However, the limited availability of clinical samples as well as the modest sample volume (~100 μL) using these collection methods presents challenges in mechanistic studies on how altered mucus composition influences its function. In addition, samples acquired from humans can often possess altered mechanical properties due to sample dilution after sputum induction [23]. *In vivo* studies in MUC5B and MUC5AC knock-out (KO) and overexpressing mouse models have provided valuable insights into their functional roles in the airways [24–26]. Specifically, it has been shown in prior work that MUC5B-deficient mice have reduced MCC, increased susceptibility to infection, and reduced survival [24]. Conversely, MUC5AC-deficient mice were protected from airway hyperreactivity upon exposure to allergens [25]. However, prior work has also shown that overexpression of MUC5AC does not negatively impact MCC and is protective against respiratory infection by influenza A virus (IAV) [26]. In addition, overexpression of MUC5B can lead to reduced MCC and contribute to lung fibrosis [25]. Thus, maintaining an optimal balance of these gel-forming mucins is likely critical to maintaining mucus function.

In our work, we report a synthetic mucus biomaterial capable of modeling transport and barrier function of airway mucus. Specifically, we have adapted an approach established in previous work [27] to create mucin-based hydrogels composed of MUC5B and MUC5AC with viscoelastic properties comparable to airway mucus. This model system provides the capability to systematically vary the composition of mucus to capture how these changes impact its physiological function. Using this model, we examined the rheology of mucin-based hydrogels with varied and specified MUC5B:MUC5AC ratio to understand the impact of mucin composition on its biophysical properties. Transport of synthetic mucus on fully differentiated human airway epithelial (HAE) tissue cultures were then compared to natively secreted mucus in normal cultures and IL-13 exposed cultures to establish an asthma-like inflammatory condition. To understand the barrier function of synthetic mucus towards respiratory viruses, we directly measured the diffusion of fluorescent IAV within synthetic mucus gels and the extent of infection in synthetic mucus-coated human airway tissue cultures. The results of work establish a new biomaterial-based approach to create *in vitro* models of asthmatic airways and identify potential mechanisms of lung dysfunction.

## 2. Methods

### 2.1 Preparation of synthetic mucus

Synthetic mucus hydrogels were prepared using a previously established method [27]. Briefly, porcine gastric mucin (PGM; Sigma-Aldrich) and mucin from bovine submaxillary glands (BSM; Sigma-Aldrich) are mixed with a cross-linking reagent, 4-arm PEG-thiol (PEG-4SH; Laysan Bio Inc.) to mediate crosslinking between mucin biopolymers. A solution of 4% (w/v) PGM and/or BSM is dissolved in a physiological buffer solution containing 154 mM NaCl, 3 mM CaCl_2_, and 15 mM NaH_2_PO_4_ at pH 7.4 at 4 % (w/v) and stirred at room temperature for 2 hours. The cross-linking reagent, PEG-4SH, is initially prepared at 4 % (w/v) in the same buffer then mixed with equal volumes of mucin solutions to achieve a final concentration of 2 % (w/v) mucin and 2 % (w/v) cross-linking reagent. For experiments with varying ratios of MUC5B and MUC5AC, solutions of PGM and BSM were mixed at varying MUC5B:MUC5AC ratios (75:25, 50:50, and 25:75) prior to addition of the cross-linking reagent. The mixed solution of mucin and cross-linker were incubated at room temperature for ≥15 hours to allow gelation. For bulk rheology experiments, synthetic mucus hydrogels were prepared in petri dishes with 40 mm diameter with 2 mm thickness. For PTM experiments, hydrogels were prepared in custom microscopy chambers prior to gelation. For mucus transport and influenza viral challenge studies, hydrogels were prepared 24 hours prior to experiments and UV sterilized for 15 minutes before being transferred to HAE tissue cultures.

### 2.2 Collection of human mucus samples

Human mucus samples were collected using the endotracheal tube (ETT) method in accordance with an IRB-approved protocol at the University of Maryland Medical Center (Protocol#: HP-00080047). ETT were collected from patients after intubation as a part of general anesthesia at UMMC. To collect the ETT-derived *ex vivo* samples, the last 10 cm of the tubes were cut, including the balloon cuff, and suspended in a 50-mL centrifuge tube using a syringe needle. The centrifuge tube containing the ETT was spun at 220 xg for 30 seconds for mucus collection, which resulted in 100-300 μL of ETT mucus. Samples were stored at −80°C immediately after collection and thawed (no more than three times) prior to use for experiments.

### 2.3 Preparation of nanoparticles for particle tracking microrheology

Fluorescent carboxylate modified fluorescent polystyrene nanoparticles (PS-COOH; Life Technologies) with a diameter of 100 and 500 nm were coated with a high surface density of polyethylene glycol (PEG) via a carboxyl-amine linkage using 5-kDa methoxy PEG-amine (Creative PEGWorks) as previously reported [21,27]. Particle size and zeta potential was measured in 10 mM NaCl at pH 7 using a NanoBrook Omni (Brookhaven Instruments). We measured diameters of 122 nm and 491 nm and zeta potentials of −0.54 ± 1.16 and 1.09 ± 0.51 mV for 100 nm and 500 nm PEG-coated PS nanoparticles, respectively.

### 2.4 Particle Tracking Microrheology (PTM) analysis in human mucus and synthetic mucus

The diffusion of the PEG-coated nanoparticles (PEG-NP) in human and synthetic mucus gels were measured using fluorescent video microscopy. Mucus samples were prepared in custom microscopy chambers that consisted of O-rings coated in vacuum grease on a microscope slide. Twenty μL of human mucus was added to the chamber along with 1 μL of ~0.002% w/v suspension of PEG-NP 30 minutes prior to PTM experiments. Synthetic mucus was prepared by adding 25 μL of mucin and cross-linker solution to the microscopy chamber. 1 μL of ~0.002% w/v suspension of PEG-NP was added to the mucin and cross-linker solution and sealed with a cover slip. Imaging was performed 24 hours later, after gelation. Fluorescent video microscopy videos were collected using a Zeiss 800 LSM microscope with a 63x water-immersion objective and an Axiocam 702 camera (Zeiss). High-speed videos were recorded at a frame rate of 33 Hz for 10 seconds at room temperature. For each sample, at least 3 high-speed videos were recorded. The particle tracking analysis was performed using a previously developed image processing algorithm [28]. Mean squared displacement (MSD) as a function of time lag (τ) was calculated as 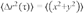, for each particle. Pore size of sample was calculated by 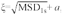, where MSD_1s_ is the measured MSD at τ= 1 second and *a* is the nanoparticle radius.

### 2.5 Measurement of bulk rheology

Dynamic rheological measurements of the synthetic mucus gels were performed using the ARES G2 rheometer (TA Instruments) with a 20-mm diameter parallel plate geometry at 25°C. To determine the linear viscoelastic region of the fully formed gel, a strain sweep measurement was collected from 0.1-10% strain at a frequency of 1 rad s^-1^. To determine the elastic modulus, *G′(ω)*, and viscous modulus, *G′(ω)* a frequency sweep measurement was conducted within the linear viscoelastic region of the gel, at 1% strain amplitude and angular frequencies from 0.1 to 100 rad s^-1^. We determined the elastic, G′ (ω), and viscous modulus, G″ (ω), by conducting a frequency sweep measurement within the linear viscoelastic region of our hydrogel, at 1% strain amplitude and angular frequencies from 0.1 to 100 rad s^-1^.

### 2.6 Culture of BCi-NS1.1 cells

The h-TERT immortalized human airway basal BCi-NS1.1 cell line was generously provided by Dr. Ronald Crystal which their group established in previous work [29]. Cells were initially seeded on plastic at ~3000 cells/cm^2^ in PneumaCult-Ex Plus media (StemCell) and incubated at 37°C, 5% CO_2_. Once cells reached 70-80% confluency, they were dissociated using 0.05% trypsin ethylenediaminetetraacetic acid (EDTA) for 5 minutes at 37°C. For differentiation at air-liquid interface (ALI), cells were seeded on 12 mm diameter transwell inserts (Corning Costar) coated with 50 μg/mL collagen from human placenta (Sigma Aldrich) at ~10,000 cells/cm^2^. Expansion media, PneumaCult-Ex Plus, was used to feed cells in both the apical and basolateral compartments until 100% confluency. After reaching confluency, the apical media was removed, introducing ALI, and the basolateral media was replaced with PneumaCult-ALI (StemCell). All cells were grown for 28 days to reach differentiation with media exchanged every other day. Collection method of BCi-NS1.1 mucus was adapted from a previously described protocol [30,31]. BCi-NS1.1 mucus was allowed to accumulate for 7 days. *In vitro* mucus samples were harvested by washing apical compartments with PBS for 30 minutes at 37°C. After 30 minutes incubation, the solution of mucus and PBS in the apical compartment was collected. Samples were loaded into Amicon Ultra 100 kDa filters (Millipore-Sigma) and centrifuged at 14,000 *xg* for 30 minutes to remove excess PBS. After filtration, mucus concentration was measured using a BCA assay and samples were reconstituted into physiological concentrations (~2 mg/mL) [32] using physiological buffer used to prepare mucin hydrogels. To establish an asthma-like human tissue culture model, BCi-NS1.1 cultures were maintained in medium with IL-13 (100 ng/mL, Peprotech, Rocky Hill, NJ) for 7 days prior to *en face* staining and MCC experiments.

### 2.7 Preparation of BCi-NS1.1 cultures for en face staining

For *en face* immunohistochemical staining and fluorescence imaging of MUC5B and MUC5AC, cultures were prepared by aspirating culture medium from the basolateral compartment and adding Carnoy solution (6:3:1 ratio of 100% ethanol, chloroform, and glacial acetic acid), a non-aqueous fixative that preserves the secreted mucus layer, to the basolateral (1 mL) and apical (0.5 mL) compartments [20]. After 30 minutes of fixation at room temperature, inserts were washed twice with absolute methanol, twice with absolute ethanol, and twice with PBS. For blocking non-specific background in all imaging experiments, inserts were incubated in 3% bovine serum albumin (BSA) in PBS, which had been filter sterilized, for 1 hour at room temperature. The primary antibodies that were used were mouse monoclonal anti-MU5AC (Santa Cruz, sc-21701, 1:100) and mouse monoclonal anti-MUC5B (Santa Cruz, sc-21768, 1:100). Primary antibody incubation was over night at 4°C in 1% BSA in PBS. After overnight incubation, inserts were washed in PBS twice and incubated in secondary antibody staining solution for 1 hour at room temperature. The secondary antibody used was Donkey anti-Mouse IgG (H+L) Highly Cross-Adsorbed Secondary Antibody, Alexa Fluor 488 (Thermo Fisher Scientific, A-21202, RRID AB_141607). Inserts were washed with PBS twice and imaged at 10x magnification using a Zeiss Axio Observer and Axiocam 503 mono camera (Zeiss). For the non-primary control, Donkey anti-Rabbit IgG (H+L) Highly Cross-Adsorbed Secondary Antibody, Alexa Fluor 647 (Thermo Fisher Scientific, catalog # A-31573, RRID AB_2536183) was used and background intensity was subtracted from anti-mucin images.

### 2.8 Measurement of mucus transport and ciliary beat frequency

Mucociliary transport was measured based on the transport of 2 μm red-fluorescent polystyrene microspheres (Sigma-Aldrich). For measurement of transport rate of native BCi-NS1.1 mucus, a 4 uL suspension of the fluorescent polystyrene microspheres (1:1000 dilution in PBS) was added on top of the native mucus, which was allowed to accumulate for 7 days. After 30 minutes of equilibration at 37°C, videos of three regions were recorded at 10x magnification using a Zeiss 800 LSM microscope. Images were collected at a frame rate of 0.5 Hz for 10 seconds on the plane of the mucus gel. Images were acquired centrally within cultures and away from the edges, where mucus tends to accumulate. For measurement of transport of synthetic mucus, BCi-NS1.1 native mucus was first removed from the cultures by washing the apical surfaces of cultures with PBS for 30 minutes. Forty μL of the hydrogel was added to the apical surface (~40 μm thickness) of fully differentiated BCi-NS1.1 cells and equilibrated for 30 minutes at 37°C. A suspension of 4 uL of 2 μm red-fluorescent polystyrene microspheres was added and allowed to equilibrate for an additional 30 minutes. For *N-acetylcysteine* (NAC) experiments, fluorescent polystyrene microspheres were diluted in PBS containing 50 mM NAC in order to prevent further dilution of hydrogels. After 30 minutes of incubation, videos of at least three regions were recorded as described for the transport rates of BCi-NS1.1 native mucus. The microsphere tracking data analysis is based on an image processing algorithm that was custom written in MATLAB (The MathWorks). Briefly, the analysis software computes the xy-plane trajectories of each fluorescent microsphere in each frame. Using the trajectory data, displacement of microspheres was computed, and transport rate was calculated by dividing the displacement of microsphere by total time elapsed. In order to measure ciliary beat frequency (CBF), 10 second videos at a frame rate of 20 Hz were recorded at 10x magnification in ≥3 randomly selected regions of HBE cell cultures using brightfield. Using a custom written algorithm in MATLAB, the number of local pixel intensity maxima were counted, which indicates beating of cilia. Beat frequency was determined by dividing the number of beats over the total elapsed time.

### 2.9 Measurement of disulfide bond concentration in synthetic mucus

Concentration of disulfide bonds were measured using a previously described fluorometric assay [19]. Synthetic mucus hydrogels were resuspended in 8 M guanidine-HCl (Sigma) to 10 times the original hydrogel volume. In order to block free cysteines, samples were treated with 10% (v/v) of 500 mM iodoacetamide for 1 hour at room temperature. To quench excess iodoacetamide and reduce existing disulfide bonds, samples were treated with 10% (v/v) of 1 M DTT at 37°C. Samples were loaded into Amicon Ultra 10 kDa filters (Millipore-Sigma) and centrifuged at 14,000 *xg* for 20 minutes for removal of excess DTT and quenched iodoacetamide. Recovered samples were resuspended in 50 mM tris-HCl (pH 8.0) and mixed with equal volumes of 2 mM monobromobimane (mBBr; Sigma). After 15 minutes of incubation in mBBr at room temperature, fluorescence was measured at 395-nm excitation/490-nm emission on a Spark Multimode Microplate reader (Tecan). Serial dilutions of 5 mM L-cysteine in 50 mM tris-HCl were used as standards. Disulfide bond concentration was calculated against the standard curve.

### 2.10 Preparation ofLAV and IAV challenge studies

The plasmid-based reverse genetics system for influenza A virus (IAV) A/Puerto Rico/8/34 (PR8; H1N1) was a gift from Dr. Adolfo Garcia-Sastre. Infectious virus was generated from cloned cDNAs in 293T and Madin-Darby canine kidney (MDCK) cell co-cultures and purified by ultracentrifugation through a 20% sucrose onto a 50% sucrose cushion as previously described [33]. The infectivity of resulting virus stocks was quantified by standard plaque assay on MDCK cells [34], yielding a viral titer of 3.9×10^9^ plaque-forming units (pfu)/mL. IAV was subsequently labeled with a lipophilic dye, 1,1’-dioctadecyl-3,3,3’3’-tetramethylindocarbocyanine perchlorate (DiI; Invitrogen). The labeled virions were then purified and concentrated via haemadsorption to chicken red blood cells [34]. IAV stock was aliquoted and stored at −80°C. This strain exhibits a primarily spherical morphology and diameter of roughly 100 nm. DiI-labeled virus stocks were counter-stained with an anti-H1N1 IAV HA antibody (NR-3148; antiserum, goat; BEI Resources, NIAID, NIH). Diffusion of IAV was assayed similarly to PTM studies. Mucin hydrogels were prepared in microscopy chambers and 1 μL of DiI-labeled IAV labeled was applied to the surface of the hydrogel and equilibrated for 30 minutes before imaging. The same tracking analysis employed in PTM experiments was used to compute the MSD of IAV. For experiments with IAV infection, BCi-NS1.1 cultures were initially washed with PBS to remove native mucus. Forty μL of hydrogels with 75:25 and 25:75 MUC5B:MUC5AC ratio were added to the apical surface and allowed to equilibrate for 30 minutes. IAV (PR8) was then applied (4 μL; 7500 plaque-forming units (pfu)) centrally to HAE cultures and incubated for 2 hours at 37°C. After inoculation, cultures were rinsed with PBS prior to a 10 minutes wash at 37°C to remove hydrogel and non-absorbed virus. Cultures were imaged 48 hours after IAV and hydrogel were removed. To image IAV infection, cultures were fixed in 1:1 methanol: acetone solution overnight at 4C. Afterwards, cultures were washed in PBS and then permeabilized in 2.5% Triton X-100 for 15 minutes. After blocking, the primary antibody that was used was mouse monoclonal anti-IAV nucleoprotein A1 and A3 blend (EMD Millipore, MAB8251, 1:250). Images were taken at 10x magnification using a Zeiss Axio Observer and Axiocam 503 mono camera (Zeiss).

### 2.11 Statistical Analysis

All graphing and statistical analyses were performed using GraphPad Prism 8 (GraphPad Software). Two-group comparisons were performed using 2-tailed Student’s *t*-test (normally distributed data) or Mann-Whitney *U* test. For comparisons between groups, one-way analysis of variance (ANOVA) followed by a Tukey *post hoc* correction was performed. Kruskal-Wallis with Dunn’s correction was used for comparison of multiple groups with non-Gaussian distributions. Bar graphs show mean and standard deviation and box and whiskers plots show median value and 5^th^ percentile up to 95^th^ percentile of the data and outliers were not included. Differences were considered statistically different at the level of *p<0.05.*

## 3. Results

### 3.1 Engineering synthetic mucus biomaterials with physicochemical properties similar to human mucus

In this study, we aimed to create a synthetic mucus biomaterial with similar rheological and transport properties to human respiratory mucus. In our previous work, we reported the design of a hydrogel system that employs a thiol-based cross-linking strategy and verified that hydrogel assembly was mediated by hydrogen and disulfide bonding [27]. Using this approach, we prepared hydrogels with 2% (w/v) mucin and 2% (w/v) cross-linking reagent, PEG-4SH, using two types of mucins: porcine gastric mucin (PGM), composed of MUC5AC [35,36], and bovine submaxillary gland mucin (BSM), composed of MUC5B [37,38] (Fig. 1A). Given the limited volume (~50-100 μL) collected *ex vivo* from patients and generated *in vitro,* we chose to use particle tracking microrheology (PTM) to characterize the samples collected to compare to our synthetic model. Our results showed that MUC5B hydrogels had similar microrheology and pore network size to HAE-derived mucus, in range of previously reported values (~200-300 nm) [21,22]. Therefore, we used PTM to compare the physical properties of our synthetic mucus biomaterial to natural human airway mucus collected *ex vivo* from endotracheal tubes (ETT) and *in vitro* collected from BCi-NS1.1 HAE cultures. Based on 100 nm PEG-NP diffusion, as measured by logarithm based 10 of mean squared displacement at time lag 1 second (log_10_MSD_1s_), we find that *ex vivo* ETT mucus had similar microstructural properties to *in vitro* mucus samples derived from BCi-NS1.1 cultures.

**Fig 1.**
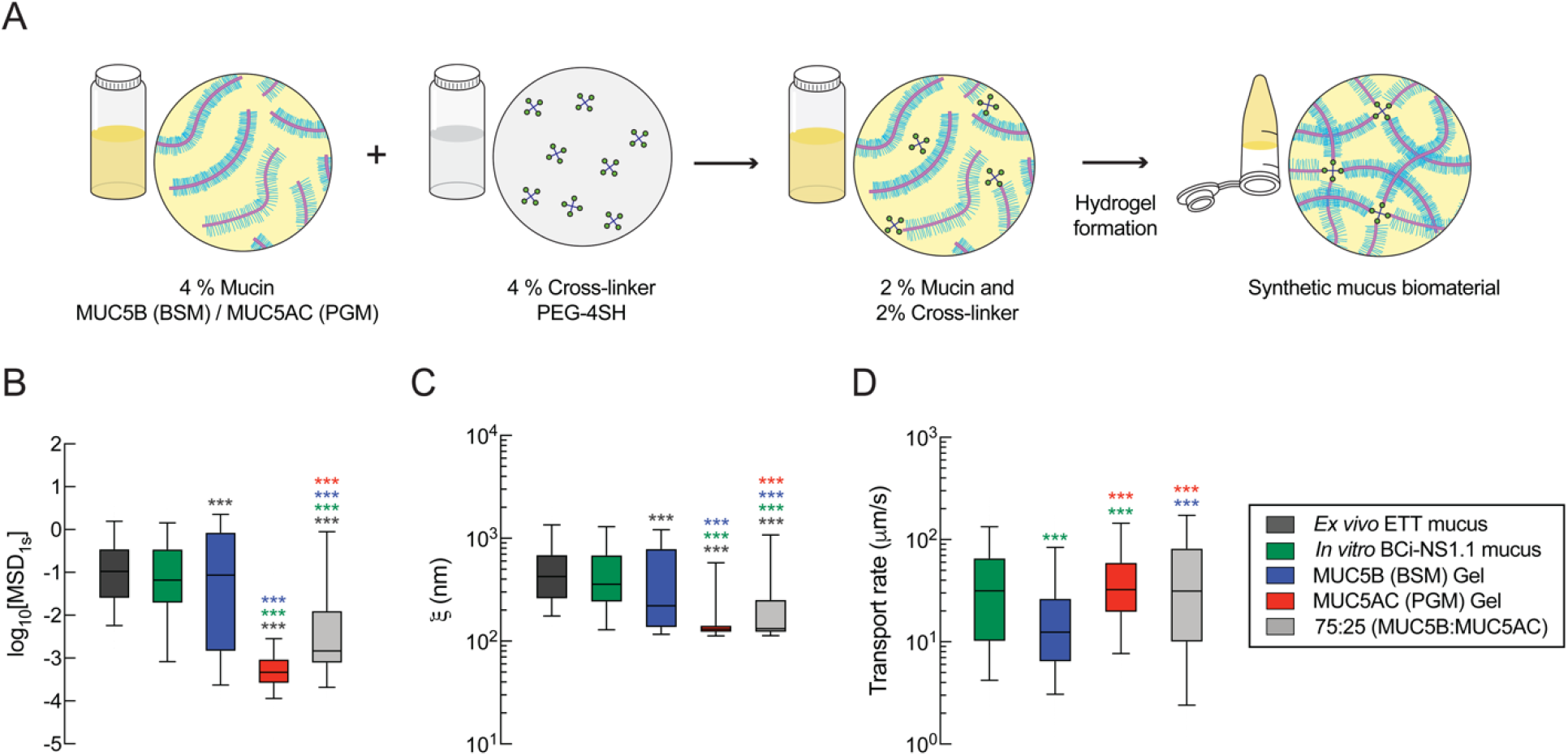
Comparison of the biophysical and transport properties of native human mucus to synthetic mucus biomaterials. (**A**) Schematic illustration of the preparation steps for the synthetic mucus biomaterial. 4% (w/v) mucin solution is mixed with 4% (w/v) cross-linker to form a mucin-based hydrogel with 2% (w/v) mucin and 2% (w/v) cross-linker. (**B**) Box-and-whisker plots of measured log_10_[MSD_1s_] for 100 nm PEG-NP in human mucus samples (*ex vivo* and *in vitro)* and synthetic mucus biomaterials with varying mucin composition. (**C**) Box-and-whisker plots of estimated pore size (ę) of human mucus samples and synthetic mucus biomaterials calculated based on 100 nm PEG-NP diffusion. (**D**) Box-and-whisker plots of microsphere transport rate of BCi-NS1.1 native mucus and synthetic mucus biomaterials on BCi-NS1.1 cultures. ***p<0.001 by Kruskal-Wallis test with Dunn’s correction. Color of asterisk indicates the comparison group.

Comparing our synthetic mucus biomaterials to human samples, we found that MUC5B (BSM) hydrogels had comparable microstructural properties to human mucus samples both collected *ex vivo* and *in vitro*(Fig. 1B). No significant differences were observed in 100 nm PEG-NP diffusion in MUC5B gels and *in vitro* BCi-NS1.1 mucus, indicating that these gels possess similar microstructural properties. However, a reduction in 100 nm PEG-NP diffusion was observed in MUC5B gels compared to *ex vivo* ETT mucus. The MUC5AC (PGM) gels had significantly reduced 100 nm PEG-NP diffusion compared to human mucus (*ex vivo* and *in vitro)* and MUC5B gels, indicative of a significantly smaller network pore size (Fig. 1B). Using PTM analysis, we estimated gel network pore sizes (ξ) in synthetic and human mucus. Similar to particle diffusion, MUC5B hydrogels showed similarities in pore size to the *in vitro* mucus samples (ξ≈200-400 nm), whereas MUC5AC gels had significantly reduced network pore sizes (ξ≈100 nm) (Fig. 1C).

Next, we measured and compared the transport rate of synthetic mucus and native mucus produced from BCi-NS1.1 HAE cultures. Although MUC5B hydrogels had similar microrheological properties to BCi-NS1.1 mucus, MUC5B gels showed significantly lower transport rates compared to native BCi-NS1.1 mucus. Alternatively, the MUC5AC hydrogel, which showed significantly reduced pore size, had significantly higher transport rates than MUC5B gels. Given native mucus contains both MUC5B and MUC5AC, we prepared a hydrogel composed of a blend of both mucins with relative concentrations representative of healthy airway mucus. Previous studies have shown that normal (non-diseased) human mucus is primarily composed of MUC5B (75-85%) and smaller amount of MUC5AC (~15-25%) [16]. To test whether a mixed ratio of mucins, specifically 75% MUC5B (BSM): 25% MUC5AC (PGM), could mimic airway mucus, we measured microrheological properties (Fig. 1B,C) and transport rates (Fig. 1D) of blended mucin hydrogels to compare to *ex vivo* and *in vitro* human mucus. Interestingly, we found the blended gels had network sizes smaller than human samples based on PTM but had the most similar transport rate to the native mucus gels produced in BCi-NS1.1 HAE cultures.

### 3.2 Macro- and microrheology of synthetic mucus with varying MUC5B:MUC5AC ratio

Previous studies have demonstrated the relative increase in the abundance of MUC5AC compared to MUC5B in asthmatic mucus from adults and children where the percentage of MUC5AC reaches up to ~70% of the total mucin content [16,39]. Therefore, we sought to determine the effects of mucin ratio, MUC5B:MU5AC, on macro- and microrheological properties of synthetic mucus biomaterials. We systematically varied the MUC5B:MUC5AC ratio starting hydrogels composed of: (i) 75% MUC5B and 25%MUC5AC, representative of health, (ii) 50% MUC5B and 50% MUC5AC representative of stable asthma and (iii) 25% MUC5B and 75% MUC5AC, representative of exacerbated asthma [16]. For simplicity, we will refer to 75:25 MUC5B:MUC5AC ratio as MUC5B-rich hydrogels and 25:75 MUC5B:MUC5AC ratio as MUC5AC-rich hydrogels. We then compared the elastic moduli of mucin-based hydrogels at ω= 1 rad s^-1^ which revealed higher elastic and viscous moduli for hydrogels with higher MUC5AC content (Fig. 2A,B). PTM analysis showed that a relative increase in MUC5AC content resulted in reduced particle diffusion, as measured by log_10_[MSD_1s_] of 100 nm PEG-NP, indicative of tightening of mesh network of the hydrogels with increased MUC5AC content (Fig. 2C). We also assessed diffusion of 500 nm PEG-NP, as PEG-NP with diameter greater than the pore size can probe physical properties closer to bulk scale [40]. The diffusion of 500 nm PEG-NP showed a systematic reduction in log_10_MSD_1s_ with increasing MUC5AC content, suggesting increased viscoelastic properties with increasing MUC5AC concentration (Fig. 2D).

**Fig. 2.**
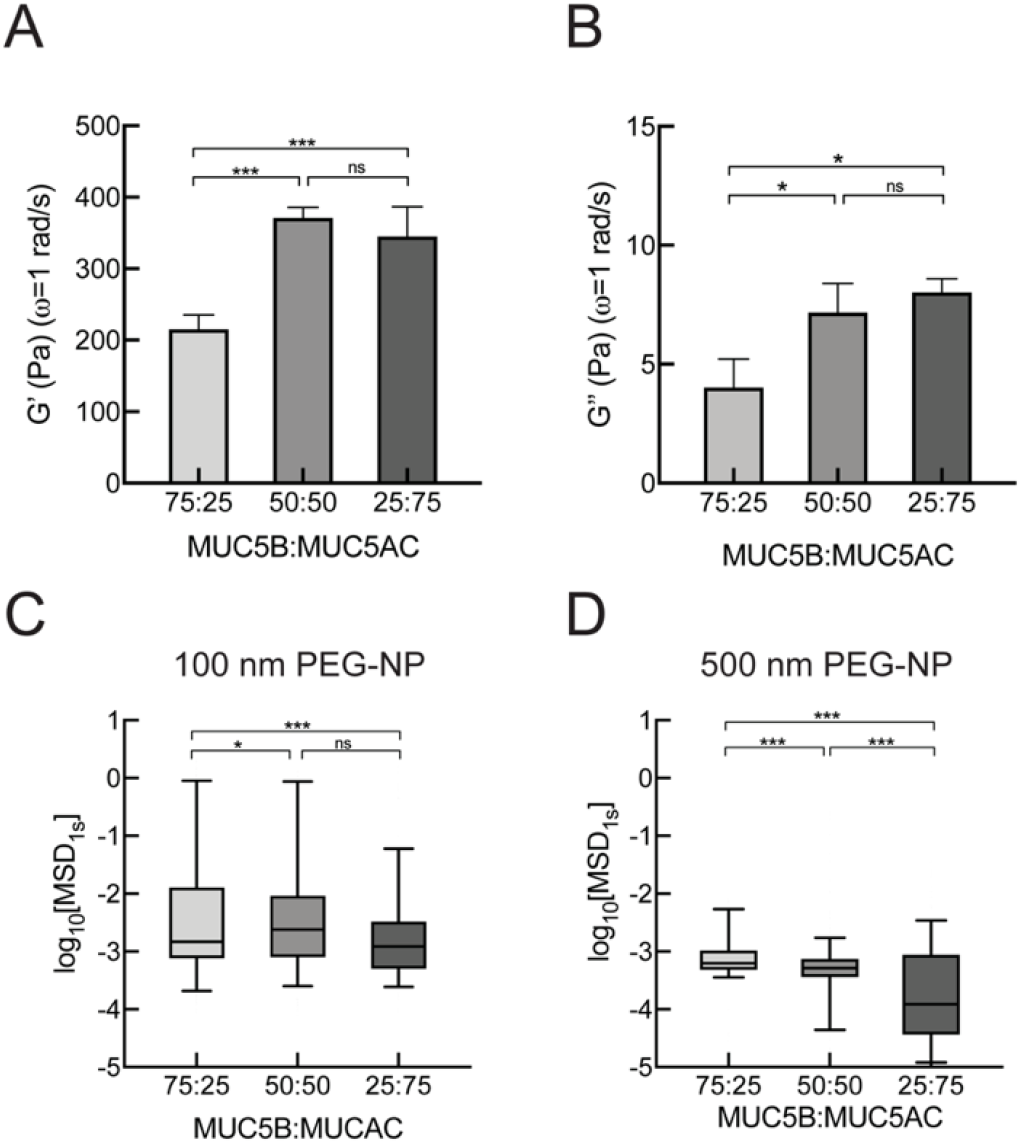
Macro- and microrheology of synthetic mucus with varied concentrations of MUC5B and MUC5AC. (**A**) Mean elastic and (**B**) viscous modulus (G’,G”) of hydrogels with varying MUC5B: MU5 AC ratio at ω= 1 rad/s measured using bulk rheology. (**C**) Box-and-whisker plots of measured logι_0_[MSD_ĭs_] for 100 nm PEG-NP in hydrogels with varying MUC5B:MU5AC ratio. (**D**) Box-and-whisker plots of measured logι_0_[MSD_ĭs_] for 500 nm PEG-NP in hydrogels with varying MUC5B:MU5AC ratio. ***p<0.001, *p<0.05 by (**A**,**B**) one-way ANOVA and (**C**,**D**) by Kruskal-Wallis test with Dunn’s correction.

### 3.3 Measurement of transport rates of synthetic mucus with varying MUC5B:MUC5AC ratio

Previous studies have shown that stimulation of primary HAE cultures with IL-13, a key mediator in asthma, results in increased MUC5AC production and markedly reduced mucus transport rates [20,41]. In order to establish and confirm asthmatic phenotype in our culture system, we stimulated BCi-NS1.1 HAE tissue cultures with IL-13 and performed *en face* staining to examine MUC5B and MUC5AC expression (Fig. 3A). IL-13 stimulation in BCi-NS1.1 cultures consistently increased MUC5AC expression, whereas MUC5B expression decreased. We observed a significant reduction in transport rates of 2 μm microspheres in IL-13 stimulated cultures (Fig. 3B,C). We also measured the cilia beat frequencies (CBF) of BCi-NS1.1 cultures to confirm that the reduction of transport rate was not due to changes in ciliary activity. We also confirmed that CBF was unaffected after IL-13 treatment (Fig. 3D). We next sought to test whether mucin composition of synthetic mucus affects mucociliary transport rates by applying hydrogels with varying MUC5B:MUC5AC ratios on apically-washed, mucus-free BCi-NS1.1 cultures. Based on microsphere trajectories, we found the transport rate of MUC5AC-rich gels were significantly reduced in comparison to gels composed of equal MUC5B/MUC5AC concentrations or MUC5B-rich gels (Fig. 3B,E). CBF was unaffected by the introduction of synthetic mucus with varied MUC5B:MUC5AC ratio (Fig. 3F).

**Fig. 3.**
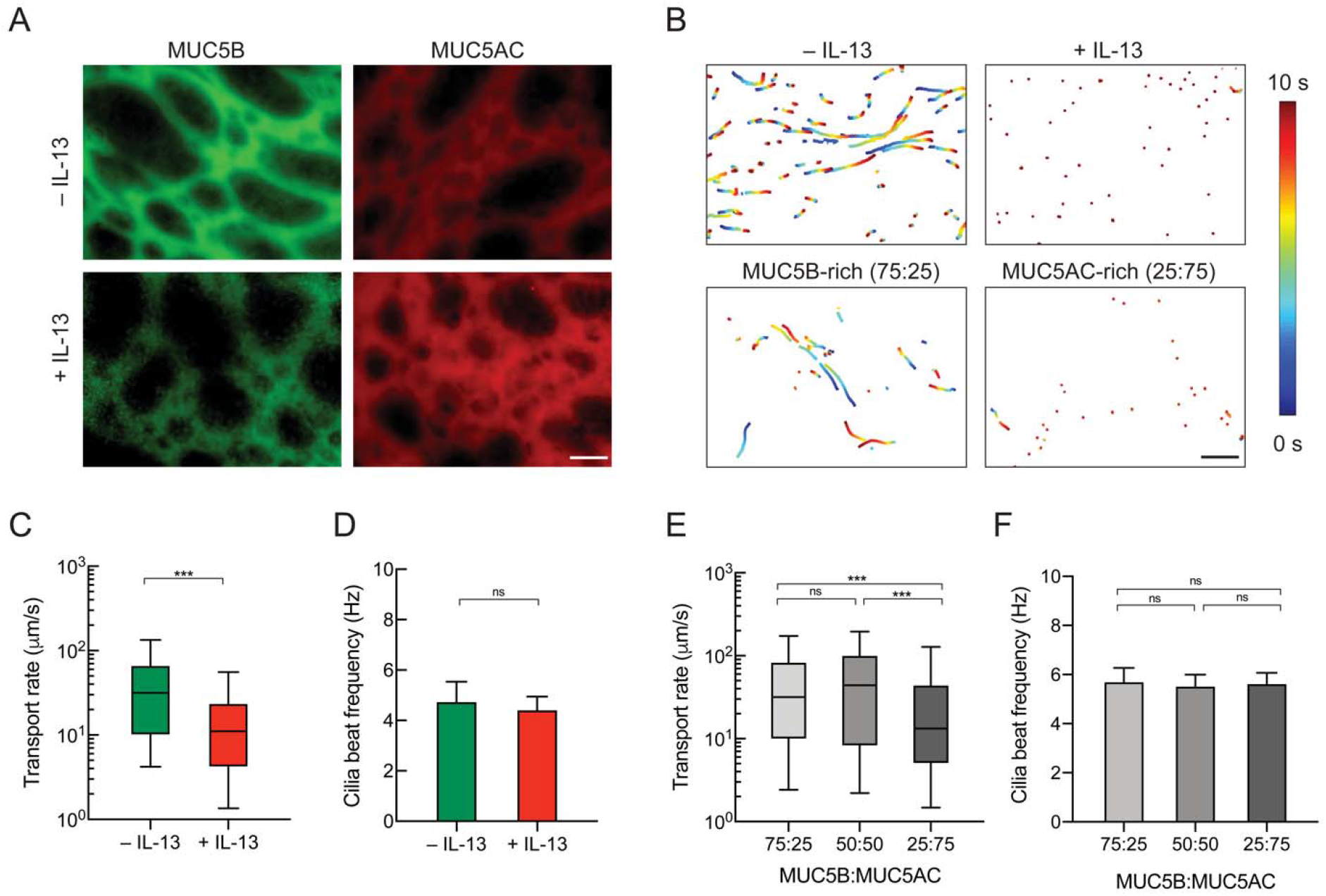
Mucociliary transport in human airway epithelial (HAE) cultures for native mucus in IL-13 stimulated HAE and synthetic mucus with varied MUC5B:MUC5AC ratio. (**A**) *En face* immunostaining of mucus gels produced from unstimulated (–IL-13) or IL-13-stimulated (+IL-13) BCi-NS1.1 HAE cultures. Scale bar = 20 μm. (**B**) Paths of fluorescent microspheres deposited onto gels from – IL-13 and +IL-13 BCi-NS1.1 cultures and synthetic mucus gels (shown conditions are MUC5B-rich (75:25) and MUC5AC-rich (25:75) gels). Trajectories show 10 seconds of motion with color scale that indicates time elapsed. Scale bar = 100 μm. (**C**) Box-and-whisker plots of microsphere transport rate on – IL-13 and +IL-13 BCi-NS1.1 cultures. ***p<0.001 by Mann-Whitney *U* test. (**D**) Mean ciliary beat frequency of – IL-13 and +IL-13 BCi-NS1.1 cultures. (**E**) Box-and-whisker plots of microsphere transport rate on BCi-NS1.1 cultures with hydrogels with varying MUC5B:MUC5AC ratio. ***p<0.001 by Kruskal-Wallis test with Dunn’s correction. (**F**) Mean ciliary beat frequency BCi-NS1.1 cultures with hydrogels with varying MUC5B:MUC5AC ratio.

### 3.4 Impact of disulfide crosslinking on the transport of asthma-like synthetic mucus gels

While a constant PEG-4SH concentration was used throughout our work, we hypothesized enhanced viscoelasticity of MUC5AC-rich synthetic mucus may be a result of the higher number of available cysteines on MUC5AC noted in a previous study [42]. Therefore, we measured the disulfide bond concentration in hydrogels that showed significant differences in macro- and microrheological properties as well as transport rates for MUC5B-rich and MUC5AC-rich gels (Fig. 4A). We found that the disulfide bond (S-S) concentration in MUC5AC-rich hydrogels was markedly higher than in MUC5B-rich hydrogels. As enhanced elasticity resulting from increased disulfide crosslinking is considered a major factor in impairing mucus clearance [19], we tested the effectiveness of N-acetylcysteine (NAC), areducing agent that is used clinically as a mucolytic agent, on restoring transport rates of MUC5AC-rich gels on BCi-NS1.1 cultures. We observed that treatment with 50 mM NAC on MUC5AC-rich hydrogels significantly increased transport rates (Fig. 4B), whereas CBF remained unchanged (Fig. 4C). While MCC was improved with NAC treatment for the asthma-like MUC5AC-rich gels, transport rates were not restored to transport values of hydrogels composed of MUC5B-rich gels, representative of health (Fig. 4B).

**Fig. 4.**
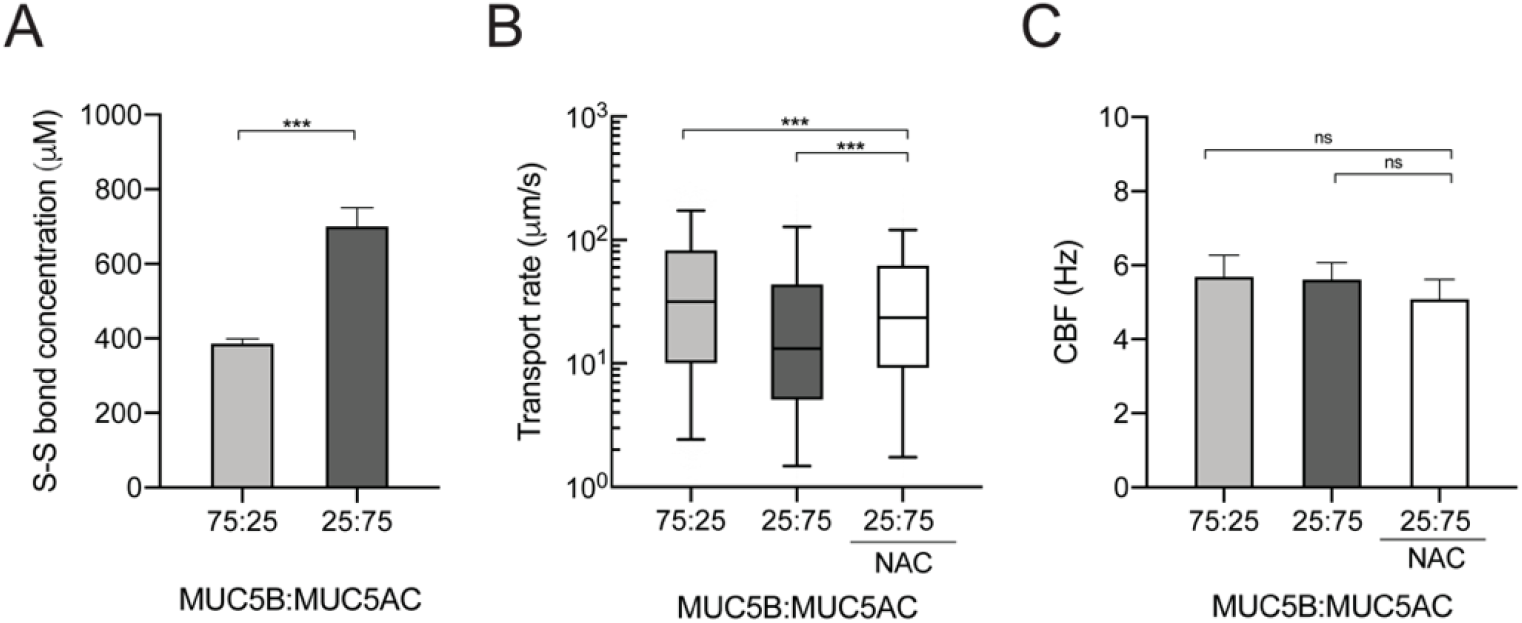
Influence of disulfide crosslinking on asthma-like synthetic mucus transport function. (**A**) Mean disulfide bond (S-S) concentration in MUC5B-rich (75:25) and MUC5AC-rich (25:75) gels. ***p<0.001 by unpaired student *t*-test. (**B**) Box-and-whisker plots of microsphere transport rates on BCi- NS1.1 HAE cultures with MUC5B-rich (75:25) and MUC5AC-rich (25:75) hydrogels and MUC5AC-rich hydrogel (25:75) with NAC treatment. ***p<0.001 by Kruskal-Wallis test with Dunn’s correction. (**C**) Ciliary beat frequency of BCi-NS1.1 HAE cultures.

### 3.5 Barrier properties of asthma-like synthetic mucus gels against influenza A viral infection

Previous studies have shown that individuals with asthma are more susceptible to viral infections [43] which can often cause an exacerbation of disease symptoms [13]. To determine if mucin composition could influence the barrier function of synthetic mucus towards influenza A virus (IAV), we measured the diffusion rate of fluorescently labeled IAV in synthetic mucus with different MUC5B:MUC5AC ratios using PTM. We found IAV had a higher average diffusion rate in MUC5AC-rich synthetic mucus compared to MUC5B-rich gels (Fig. 5A,B). However, the frequency distribution of log_10_[MSD_1s_] of individual IAV showed that only a small fraction of IAV could rapidly penetrate MUC5B-rich (~7%) and MUC5AC-rich gels (~9%) (Fig. 5C). We further tested if synthetic mucus with variable mucin content were more or less effective at blocking IAV infection in BCi-NS1.1 HAE cultures. In these studies, BCi- NS1.1 cultures were coated with MUC5B-rich (75:25) and MUC5AC-rich (25:75) synthetic mucus and challenged with 7500 pfu of IAV (multiplicity of infection ~ 0.15). IAV were introduced in minimal volume (4 μL) in order to avoid diluting mucus and the inoculum were kept on the cultures for 2 hours. Two days post infection, staining for IAV nucleoprotein revealed a significant reduction in successful infection by IAV when coated with MUC5B-rich synthetic mucus whereas MUC5AC-rich, asthma-like mucus had a similar amount of staining for IAV nucleoprotein to uncoated controls.

**Fig. 5.**
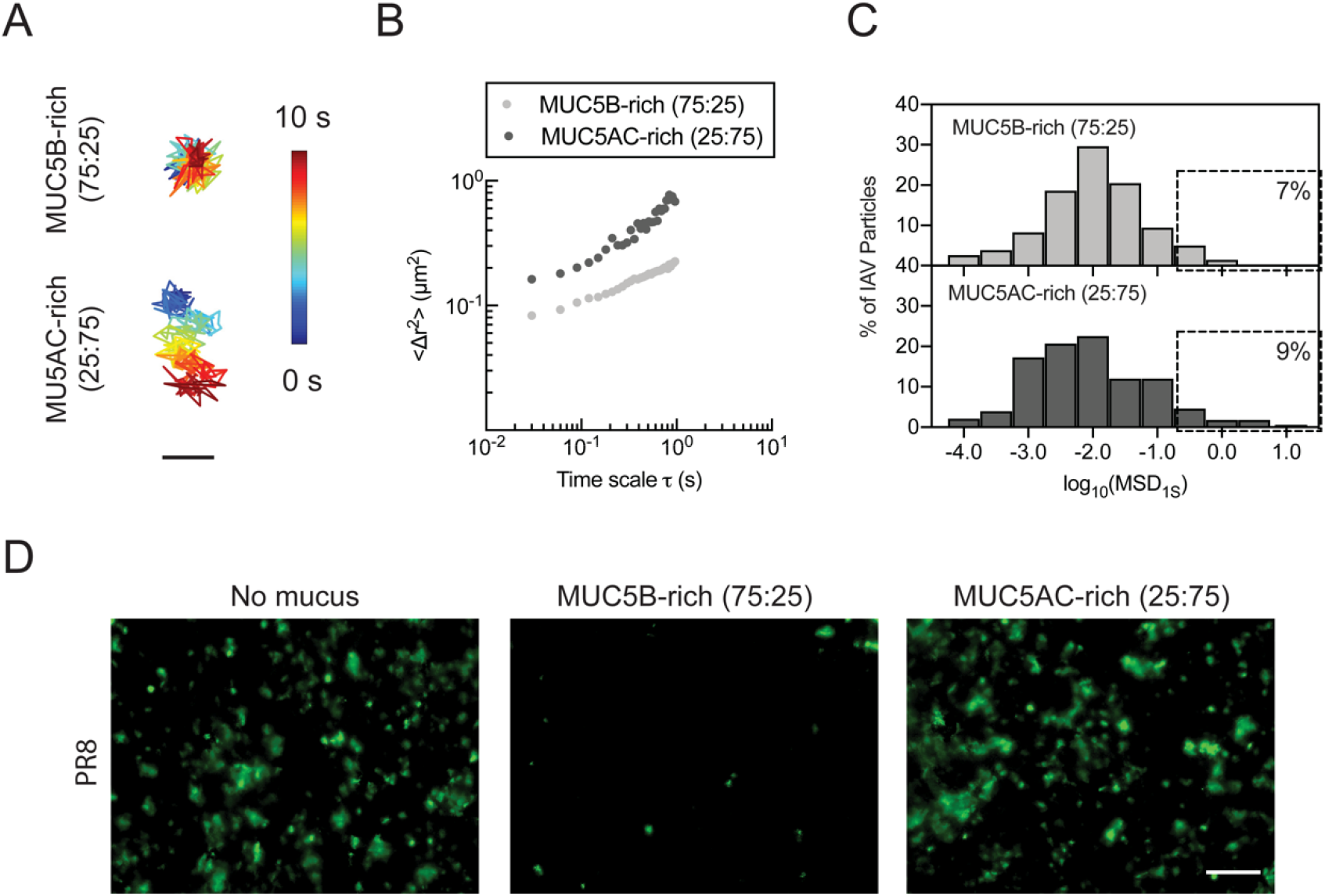
Barrier function of normal and asthma-like synthetic mucus towards influenza A virus (IAV). (**A**) Trajectories of IAV diffusion in MUC5B-rich (75:25) and MUC5AC-rich (25:75) gels. Trajectories show 10 seconds of motion with color scale that indicates time elapsed. Scale bar = 200 nm. (**B**) Ensemble average MSD (<△r^2^>) of DiI labeled IAV as a function of time scale (τ) in MUC5B-rich (75:25) and MUC5AC-rich (25:75) gels. (**C**) Frequency distribution of log_10_[MSD_1s_] of individual IAV in MUC5B-rich (75:25) and MUC5AC-rich gels (25:75). (**D**) Fluorescent micrographs of uncoated and synthetic mucus coated BCi-NS1.1 HAE cultures infected apically with IAV (PR8) 48hpi after 2 hour inoculation. Green indicates staining for IAV nucleoprotein. Scale bar = 20 μm.

## 4. Discussion

In order to create synthetic mucus biomaterials with native-like function, we formulated mucin hydrogels using BSM, primarily composed of MUC5B, and PGM, primarily composed of MUC5AC, combined with a star PEG-thiol cross-linking agent to establish rheological properties similar to human airway mucus. We directly compared our synthetic mucus model to human mucus collected (i) from individuals without lung disease and (ii) from well-differentiated HAE tissue cultures. Although MUC5B gels were most similar to HAE-derived mucus, we observed significant differences between MUC5B gels and ETT- sourced mucus. We expect these differences to arise due to heterogeneity in microrheological properties in clinical samples of mucus as these samples possess high amounts of non-mucin biomolecules that can contribute to gel organization and microstructure [23]. MUC5AC hydrogels, on the other hand, had significantly smaller pore sizes (~100 nm) compared to ‘normal’*ex vivo* and *in vitro* human mucus that were comparable to previously reported properties of mucus in diseases, such as in CF [22]. We measured total protein content as well as hydrodynamic radii of mucin solutions and confirmed no significant differences between protein concentration and size of MUC5B (BSM) and MUC5AC (PGM) solutions (**Fig. S1**).

Due to the similarities in microrheological properties between MUC5B gels and HAE-derived mucus, we expected comparable transport rates on HAE cultures. Interestingly, transport rates of MUC5B gels were significantly lower compared to native HAE mucus (Fig. 1), whereas MUC5AC gels exhibited significantly faster transport rates compared to MUC5B hydrogel, despite differences in microrheological properties. Thus, neither MUC5AC or MUC5B hydrogels alone resembled both the biophysical and transport functions of native human mucus, which may suggest MUC5B transport could be influenced by factors other than network elasticity, such as friction or surface adhesion [44,45]. As native airway mucus is composed of both MUC5B and MUC5AC, we prepared hydrogels with both mucins at a physiological MUC5B:MUC5AC ratio of 75:25 (MUC5B-rich). Our results showed that blending of the mucins resulted in mucus clearance rates that were comparable to the transport of BCi-NS1.1 HAE mucus and also in range of physiological values reported in literature (~50 μm/s) [45]. Our results would suggest both mucins work in concert to enable efficient mucus transport at airway surfaces. We should note the bulk elastic moduli measured here, on the order of 200-400 Pa, are higher than previously reported moduli for human airway mucus (~10-50 Pa) [18,23,30]. With the similarities in transport rates of synthetic and native human mucus on airway surfaces (Fig. 1E), we expect synthetic mucus is partially diluted and as a result, its elasticity is reduced when equilibrated for 30 minutes on HAE tissue culture prior to experiments.

Our results suggest, independent of concentration, increases in MUC5AC, as observed in sputum samples from patients with asthma, could be sufficient to enhance the bulk viscoelastic properties of mucus (Fig. 2). In the past, these changes in viscoelastic properties in asthma have been attributed to increased mucin concentration and excess plasma protein that may interfere with normal degradation and turnover of mucins in the airways [18]. While we presume transport in IL-13 stimulated HAE will also be influenced by MUC5AC tethering to HAE as previously shown [20], our data on synthetic mucus transport suggests increasing MUC5AC in mucus gels may progressively slow MCC (Fig. 3). We also found the enhanced elasticity and reduced transport of MUC5AC-rich gels is likely due to increased mucin-mucin disulfide cross-linking (Fig. 4). Given the importance of disulfide bonds in mucus gel elasticity, reducing agents such as N-acetylcysteine (NAC) have been used as an inhaled treatment to breakdown hyper-viscous mucus in chronic lung diseases like cystic fibrosis (CF) and chronic obstructive pulmonary disease (COPD) [25,46]. Using NAC, we also found reducing the disulfide bonds can significantly improve transport of asthma-like synthetic mucus with increased MUC5AC (Fig. 4B).

Given susceptibility to infection in those with underlying diseases like asthma [12], we examined the role of MUC5B:MUC5AC ratio in the barrier function of synthetic mucus towards IAV. We expected the tighter microstructure of MUC5AC-rich gels to reinforce barrier properties against IAV, restricting IAV diffusion. In contrast, we observed faster average diffusion of IAV in asthma-like, MUC5AC-rich gels compared to MUC5B-rich gels (Fig. 5B). However, in both MUC5B and MUC5AC-rich gels, we estimate only a small fraction (7-9%) of IAV particles can diffuse rapidly enough to penetrate a ~40 μm mucus layer during the 2-hour period of exposure to viral inoculum used in our studies (Fig. 5C). While these gel thicknesses and exposure times are more reflective of diseased airways, we note thinner mucus layers and shorter clearance times (on the order of 30 minutes) would be expected in healthy lungs *in vivo*[45]. However, we kept gel thickness and inoculation time consistent for both MUC5B and MUC5AC-rich to enable direct comparisons. Notably, we found in HAE cultures challenged with IAV that the barrier properties of MUC5AC-rich gels towards IAV are significantly compromised whereas MUC5B-rich gels are effective in neutralizing IAV (Fig. 5D). It has been established in prior studies that IAV interacts with mucin glycans, primarily sialic acid [47]. In comparing mucin glycosylation profiles, MUC5B has been shown to contain a high concentration of sialic acid whereas MUC5AC contains greater fucose content [3,48]. Thus, we expect that the reduction in sialic acid content and transport rates of MUC5AC-rich gels could render the airway surface more susceptible to widespread and increased frequency of infection. We anticipate that the barrier properties of MUC5B and MUC5AC can be further understood by studying IAV diffusion through and extent of infection in synthetic mucus-coated cultures using gels with modified glycosylated domains. Modification of mucin glycans, specifically removal of sialic acids, of BSM gels through enzymatic cleavage using sialidase has been demonstrated in prior work [49] and will be pursued in our future studies.

## 5. Conclusion

Our results establish a new biomaterial capable of recapitulating native airway mucus properties where mucus composition can be precisely controlled to study how its functions may be altered in disease. Using bioengineered synthetic mucus, we have found that the changes observed in mucus composition in asthma can have a major impact on gel viscoelasticity which in turn reduces its capacity to be effectively cleared. We also observed asthma-like mucus biomaterials were less capable of protecting the airway surface from infection by IAV. This work motivates future studies on patient-derived human mucus from individuals with asthma to determine how rheology and/or barrier function towards respiratory viruses may be affected by changes to mucin composition. The model established herein could also be applied to other lung diseases (e.g. COPD, CF, pulmonary fibrosis) to understand airway dysfunction and uncover new disease mechanisms that provide the basis for development of diagnostic biomarkers and therapeutic interventions.

## Statement of Significance

Clinical studies have shown mucus obtained from individuals with asthma possesses altered mucin composition. However, how these changes alter the functional properties of airway mucus is not well understood. To address this, we have developed a synthetic mucus biomaterial that mimics the properties of native airway mucus. We demonstrate using synthetic mucus biomaterials how the changes in mucin composition observed in asthma may cause the mucus gel to become immobile which in turn reduces its ability to be cleared and increases the likelihood for mucus plug formation in the lungs. In addition, we demonstrate how the barrier function of asthma-like synthetic mucus towards influenza A virus is impaired and may lead to worsening of respiratory infection.

## Declaration of Competing Interest

The authors declare no conflict of interest.

## Acknowledgements

This work was supported by the University of Maryland, Burroughs Wellcome Fund Career Award at the Scientific Interface (to G.A.D.), Parker B. Francis Fellowship in Pulmonary Research (to M.A.S.), Cystic Fibrosis Foundation (DUNCAN18I0 to G.A.D. & M.A.S.), and the NIH (R21AI142050 to M.A.S.& G.A.D., T32AI125186A to E.B.I).

## Notes

### Competing Interest Statement

The authors have declared no competing interest.

