## Supplemental Information for "Modeling airway dysfunction in asthma using synthetic mucus biomaterials"

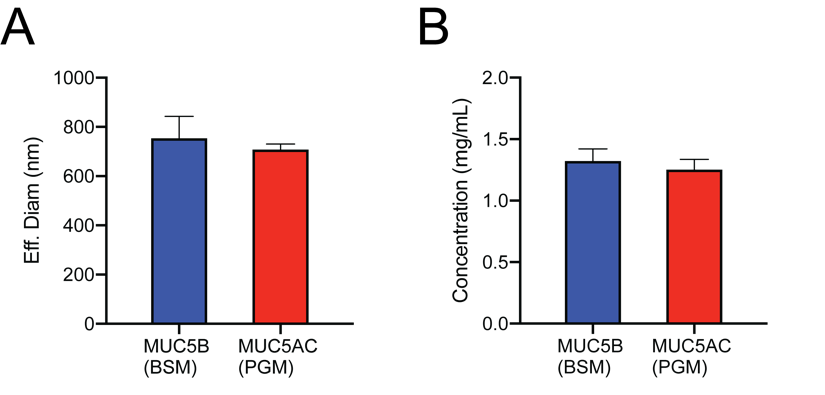

**Fig. S1**. (A) Effective hydrodynamic diameter of MUC5B (BSM) and MUC5AC (PGM) solutions measured in 10 mM NaCl using dynamic light scattering (NanoBrook Omni; Brookhaven Instruments). (B) Concentration of 1% mucin solutions measured using BCA assay.
